# Taxonomic filtering accompanies functional expansion during long-term soil restoration

**DOI:** 10.64898/2026.03.17.712278

**Authors:** Tim Goodall, Susheel Bhanu Busi, Briony Jones, Amy Thorpe, Robert I. Griffiths, John Redhead, Lucy Hulmes, Sarah Hulmes, Lucy Ridding, Jodey Peyton, Gloria Pereira, Hyun Soon Gweon, Daniel S. Read, Richard Pywell

## Abstract

The restoration of species-rich calcareous grasslands is a critical conservation objective, yet the recovery of the invisible below-ground microbiome remains poorly quantified compared to above-ground vegetation. Using a unique 143-year land-use chronosequence on Salisbury Plain, UK, we investigated the trajectory of ecosystem reassembly across arable, regenerating (23 and 67 years), and ancient grasslands. By integrating vegetation surveys with soil physiochemistry, microbial profiling, and shotgun metagenomics, we identified a profound functional decoupling between floral and edaphic recovery. While vegetation diversity recovered relatively rapidly, approaching saturation within 23–67 years, soil properties exhibited persistent legacy effects and slow convergence. Bacterial richness decreased with restoration age, reflecting a transition from disturbance-adapted copiotrophs in arable soils to a specialised, oligotrophic community in ancient sites. This taxonomic contraction was conversely matched by an expansion in functional potential, driven by the emergence of specific taxa (e.g., Microthrixaceae, *Aquihabitans* sp.) and metabolic pathways associated with complex carbon cycling and stress tolerance. Crucially, the soil ecosystem did not reach equilibrium even after 67 years, characterised by persistent legacy phosphorus and a slow accumulation of soil organic matter. These findings suggest that passive regeneration alone may be insufficient for full soil functional recovery, and that strategies targeting microbial assembly and long-term carbon dynamics warrant further evaluation.

## Introduction

Calcareous grassland are among the most species-rich ecosystems in Northern Europe, supporting exceptional floral diversity and providing critical ecosystem services, including carbon sequestration and hydrological regulation^1^. However, the extent and quality of these habitats have declined precipitously over the last century due to agricultural expansion and intensification^2^ This conversion to agricultural land, typically involving deep tillage and the application of inorganic fertilisers, fundamentally alters the soil’s physicochemical architecture and disrupts the complex biotic interactions of the below-ground microbiome^3^. While the immediate impacts of agricultural intensification are well-documented^4^, the capacity for these ecosystems to recover following the cessation of intensive agricultural practice, i.e. natural regeneration^5^ or passive restoration^6^, remains a subject of significant ecological debate^7^.

A central uncertainty in restoration ecology is the degree of hysteresis present in soil systems. While above-ground vegetation communities may begin to visibly reassemble once disturbance pressures are removed^8^, soil ecosystems often exhibit a legacy effect^9,10^. Anthropogenic inputs, particularly elevated levels of phosphorous and nitrogen, can persist for decades^11^, potentially locking the ecosystem into an alternative trajectory, inhibiting the return of characteristic grassland biodiversity^12^. Furthermore, it is unclear whether the recovery of soil microbial diversity and function tracks linearly with vegetation recovery, or if it operates on a decoupled, much slower trajectory. Understanding these temporal dynamics is critical, as microorganisms mediate key biogeochemical cycles that underpin the long-term stability of the restored ecosystem.

Our study investigates these dynamics using a unique 143-year chronosequence located on Salisbury Plain in the United Kingdom^10^. This landscape serves as a rare ‘space-for-time’ substitution experiment, where historical land-use records allow us to identify areas of calcareous grassland with distinct cessation dates of agricultural practices. By analysing soils from active arable land (0 years) and comparing them to grasslands regenerating for 23 years (recent), 67 years (middle), and over 143 years (old/ancient), we can quantify the rate and direction of edaphic functional change. Specifically, we aimed to address three critical knowledge gaps: 1) Does the invisible below-ground ecosystem recover at the same rate as the visible above-ground flora? 2) Do soil nutrient stoichiometries and microbial community structures ever reach a stable equilibrium, or do they continue to evolve over generational-scale timeframes? 3) Which specific microbial taxa and metabolic functions act as indicators of successful restoration? By integrating vegetation surveys with detailed edaphic analysis, microbial community profiling, and metagenomic sequencing, our research provides a comprehensive view of ecosystem recovery at a landscape scale.

## Methods

### Study site and land history

The Salisbury Plain site falls under the management of the UK Ministry of Defence and represents an area of ca. 38,000 ha. As a result of its use as a military training area for over a century, much of the landscape has remained inaccessible for agriculture beyond extensive livestock grazing and thus has not been subject to the agricultural intensification which destroyed calcareous grasslands across much of Western Europe over the twentieth century. Salisbury Plain thus contains Western Europe’s largest remaining fragment of lowland calcareous grassland and forms a unique landscape for exploring spatiotemporal patterns and processes in this habitat. Approximately 37% of the current military training area consists of high-quality, species rich calcareous grassland communities, with the remainder being a mix of mesotrophic grassland, agriculturally improved pastures and arable land. The land history classifications determined by Redhead et al. (2014) from a time series of historic maps^10^, were used to determine sites suitable for analysis in this study. In total 7 arable, 6 recent, 5 middle and 7 ancient sites were chosen

The minimum age of unimproved grassland are as follows: a) Arable, 000 years denotes areas lost to agricultural improvement and currently manged for arable crops, b) Recent, 023 years; grassland from 1985 to 1996, c) Middle, 067 years; grassland from 1930 to 1967, and d) Ancient, 143 years; grassland from 1840 to 1880

Soil samples were collected in January 2013. Per site one sample was derived from the homogenisation of five sub-samples, collected along a 100 m linear transect using 5 cm virgin-plastic corers inserted into the soil to a depth of 15 cm (unless encountering an impenetrable chalk horizon). From each transect, soil from below the organic horizon was homogenised and stored in clean plastic bags and transfer to the laboratory for subsequent processing and storage at –20 °C.

### Vegetation survey

Vegetation surveys were conducted in summer 2013^10^. Briefly, for each discrete classification (arable, recent, middle and ancient), the cover of vascular plant species was recorded from five square quadrats (each 2 m x 2 m) placed at 20 m intervals along the same 100 m transect as used for soil sampling. Cover of all vascular plant species was recorded using the DAFOR scale^13^. DAFOR values used for Shannon’s calculations were Dominant = 5, Abundant = 4, Frequent = 3, Occasional = 2, Rare = 1, Present = 0.1.

### Soil physicochemical properties

Subsamples of soil were sent to the commercial soil analysis provider NRM-Cawood Scientific, Bracknell, UK and used to determine soil pH, total and organic carbon (% dry soil), phosphorus (mg / kg dry soil), potassium (mg / kg dry soil), magnesium (mg / kg dry soil), and soil texture percentages (clay <0.002 mm, silt 0.002-0.05 mm, sand 0.05-2.00 mm). Analysis of soil moisture (%), organic matter by loss on ignition (LOI % dry soil), total and organic nitrogen (% dry soil) were undertaken at the UKCEH Centralised Chemistry laboratories (Lancaster) using established protocols^14^. C:N ratios were calculated from organic carbon and nitrogen values.

### PLFA analysis

Phospholipids were extracted from 1.5 g soil fresh weight as per Whitaker *et al.* (2014) ^15^. The following Fungal and Bacterial markers were summed for each sample and the fungal to bacteria ratio calculated. Fungal markers - C18:1ω9c, C18:2ω6c, C18:2ω9t. Bacterial markers (Gram+ branched, Gram-monoenoics, cyclopropyl, actinomycetes) – “i-C15:0”, “a-C15:0”, “i-C:16:0”, “i- C17:0”, “a-C17:0”, “i-C18:0”, “a-C18:0”, “C16:1ω7c”, “C16:1ω9c”, “C17:1ω10c”, “C18:1ω11c”, “C18:1ω11t”, “cy-C17:0*”, “9,10-cy-C19:0”, “11,12-cy-C19:0”, “10Me-C16:0”, “10Me-C17:0*”, “10Me-C18:0*”.

### DNA extraction

Each sample was defrosted and DNA was extracted from 0.2 g of sample using the Mobio PowerSoil DNA extraction kit (Mobio, US) following manufacturer’s instructions. Briefly, samples were lysed at 25 Hz for 20 minutes using a TissueLyser II (Qiagen, Germany). Purified DNA was eluted in 100 µL of elution buffer. DNA purity was assessed using a NanoDrop 8000 spectrophotometer (Thermo Fisher, UK), and concentration was measured using the Qubit dsDNA kit (Thermo Fisher, UK). DNA was archived at −20 °C at UKCEH, Wallingford prior to amplicon and metagenomics analysis.

### Metagenomic sequencing and data processing

Library preparation and sequencing were conducted by Mr DNA (Texas, US). Samples underwent 2x150 bp shotgun metagenomic sequencing on an Illumina HiSeq, targeting a depth of at least 4 Gb raw data per sample. The process generated a mean number of 9.8e+06 raw reads per sample.

Data processing followed analyses pipelines implemented using the Snakemake^16^ workflow management system v7.8.2. Illumina adaptor sequences were trimmed, and reads were filtered to a minimum quality score of 25 using Trim Galore v0.6.5. Reads mapping to the human reference genome (GRCh38) were removed. To profile the community composition, SingleM v0.16.0^17^ was used to determine the percentage of archaea, bacteria, and non-prokaryotes which includes fungi and other eukaryotes. Taxonomic composition was profiled using SingleM within a reproducible workflow implemented in Snakemake. For each sample, paired-end reads were analysed with *singlem pipe* against the curated SingleM metapackage, which was downloaded automatically using *singlem data* to ensure consistent reference versions across runs. Outputs from all samples were subsequently integrated using *singlem summarise* to produce combined abundance matrices and species-by-site relative abundance estimates. The fraction of metagenomic reads derived from prokaryotic cells (Bacteria and Archaea), hereafter termed the SingleM prokaryotic fraction (SPF), was estimated using singlem prokaryotic_fraction.

### Assembly and functional annotation

Reads were assembled into contigs using Megahit v1.2.9^18^, which served as the basis for downstream annotation and analysis. For functional analysis, Open Reading Frames (ORFs) were predicted using Prodigal v2.6.3^19^ and annotated using EggNOG-mapper v2.1.9^20^ against the v6.0 database and the SEED classification database^21^. Gene abundance was calculated using featureCounts^22^. Taxonomic annotation of contigs was performed using Kaiju^23^ using a 0.7 confidence threshold against the ‘kaiju_db_nr_euk’ database (https://bioinformatics-centre.github.io/kaiju/downloads.html), and Bracken v2.6.0^24^ against the PlusPFP database^25^.

### Invertebrate isolation

Samples for extraction of soil invertebrates were collected in October 2015. Four 10 cm diameter cores were taken at each site by placing a plastic ring on the surface and using a knife to cut the surface vegetation and subsurface roots. The turf and first 10cm of soil were then lifted out as a plug and placed in clean plastic bags. Within 24 hours, these plugs were positioned in Tullgren funnels, and all invertebrates leaving the sample were captured and stored in 100% ethanol. The invertebrate catch were subsequently removed from the storage ethanol, dried and placed into Mobio PowerSoil DNA extraction kit (Mobio, US) for DNA extraction following manufacturer’s instructions.

### Amplicon sequencing

Amplicon libraries were constructed according to the dual indexing strategy of Kozich et al., 2013^26^ with each primer consisting of the appropriate Illumina adapter, 8-nt index sequence, a 10-nt pad sequence, a 2-nt linker and the amplicon specific primer. For the V3-V4 region of 16S rRNA (bacteria) CCTACGGGAGGCAGCAG and GGACTACHVGGGTWTCTAAT^26^, ITS2 (fungi) GTGARTCATCGAATCTTTG and TCCTCCGCTTATTGATATGC^27^ and COI (invertebrates, predominantly Arthropoda, Mollusca and Nematoda) GGWACWGGWTGAACWGTWTAYCCYCC and TAIACYTCIGGRTGICCRAARAAYCA^28^. The pooled libraries were sequenced separately on Illumina MiSeq (Illumina, US) V3-600 cycle flow cells and demultiplexed at UKCEH. Sequence tables for 16S and COI amplicons were generated by the process outlined by Jones *et al.* (2021)^29^, the ITS fungal amplicon sequence table was produced using PIPITS^30^. After initial quality checks using R package *microeco*^31^ sample reads were rarefied to the minimum sample read number per amplicon of 2385, 4792 and 8184 for COI, 16S and ITS reads respectively.

### Statistical analysis

Statistical analyses assessed the recovery of soil biotic properties across the restoration chronosequence. R package *vegan*^32^ produced alpha diversity metrics, including Shannon’s diversity index, Species Richness, and Pielou’s evenness, to quantify structure. The co-variance of measured edaphic variables were assessed using Pearsons and hierarchical clustering, subsequently soil organic matter (LOI), C:N, organic nitrogen, magnesium, phosphorous, potassium, pH, moisture and clay were used in the following analysis. Functional compositional differences between restoration ages were visualized using Principal Component Analysis (PCA) with vectors of edaphic drivers added. Functional annotations across hierarchy levels (SEED levels 1–4) were reshaped into long format and summarised by restoration age. For each function and time point, mean relative abundance was calculated across samples. To identify functions exhibiting consistent directional change over restoration, mean relative abundances were compared across the four time points (000, 023, 067, and 143 years). Functions were classified as emergent if they were absent at baseline (mean abundance = 0 at 000 years) and exhibited monotonic increases thereafter, and as declining if they were present at baseline and exhibited monotonic decreases across successive time points, with a net decrease from 000 to 143 years. For each function, the magnitude of change was quantified as the difference in mean relative abundance between 143 and 000 years. For visualization, functions were grouped by SEED hierarchy level and direction of change (emergent or declining), and the top functions within each group were selected based on the absolute magnitude of change. Mean relative abundances at each time point were displayed as a heatmap, with restoration age on the x-axis and functions on the y-axis, ordered by magnitude of change within each ontology level. Tile colour represents mean relative abundance and was log₁₀-transformed to enhance contrast among low-abundance functions. This approach emphasizes both the temporal dynamics and ontological structure of functional shifts during ecosystem restoration. Direct relationships between functions and soil organic matter (LOI) were assessed using Spearman’s rank correlation coefficient.

### Data and code availability

Scripts and data tables are available at https://doi.org/10.5281/zenodo.18632861

Sequence data is available at NCBI-SRA under BioProject ID: PRJNA1424699

## Results

### Response speeds and lack of equilibrium

To understand how the passive restoration of these soils might influence various soil and vegetative properties, we assessed several factors over the course of the recovery period (Figure 1A-1D). Analysis of the rates of change (Figure 1A and 1B) demonstrate that restoration is a non-linear process. The highest rates of change occurred in the initial, 000-to-023-year, interval, driven by very large increases in vegetation richness and decreases in soil phosphorus. However, the system did not stabilise after this initial burst. Between 067 and 143 years, significant changes continued to occur, particularly in soil organic matter (LOI) accumulation, moisture retention, and magnesium levels (Figure 1B and 1D). This indicates that even after nearly seven decades of recovery, the soil ecosystem had not yet reached the functional equilibrium observed in ancient grasslands.

**Figure 1.**
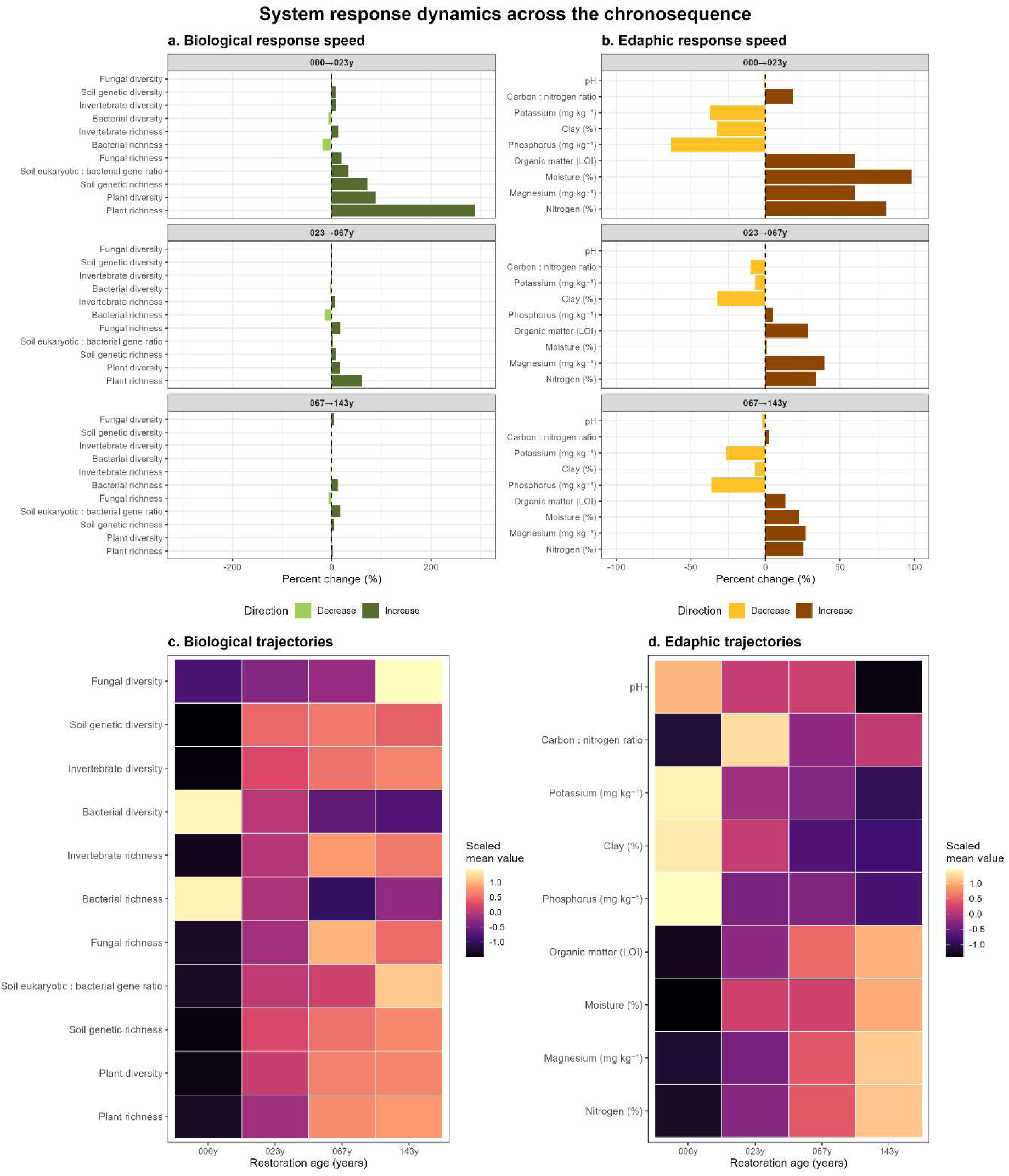
System response dynamics across a restoration chronosequence. Response speed of biological (a) and edaphic (b) variables shown as percent change between successive restoration intervals (0–23, 23–67, and 67–143 years), and standardized trajectories of the same variables across the full chronosequence (c–d). Bars indicate the magnitude and direction of change within each interval, while heatmaps illustrate relative temporal trajectories across restoration ages. Variables are ordered by the absolute magnitude of percent change across the chronosequence to facilitate comparison of response strength and temporal dynamics.

### Biological and edaphic recovery is decoupled

The restoration trajectories revealed a decoupling between plant community reassembly and soil chemical recovery (Figure 1A and 1B). Soil chemical variables exhibited a slower, continuous recovery, where soil organic matter (LOI) and nitrogen (N) concentration increased monotonically from the arable baseline to the 143-year ancient grasslands (Figure 1D). Whilst legacies of agricultural fertilisation persisted with available phosphorous (P) and potassium (K) remaining elevated in younger restoration sites compared to ancient grasslands, declining gradually over time.

Vegetation richness and diversity (Shannon index) responded more rapidly than soil chemical properties after agricultural cessation, increasing significantly in the first 23 years (surging by nearly 300%), a further increase in richness occurs 023 to 067 years and then gradual, but non-statistically significant, increases occur between 067-to-143-year stage (Supplementary Figure 1).

### Microbial variance over time

The recovery of the soil microbiome followed a trajectory distinct from that of the vegetative community (Figure 1A and 1C), characterised by an inverse relationship between taxonomic diversity and functional potential. Contrary to the trajectory observed in plant communities, soil bacterial diversity decreased with restoration age. Both bacterial richness and diversity were significantly higher in arable soils (000 years). The cessation of agricultural practices resulted in a sharp decline in bacterial richness during the initial restoration phase, with richness remaining lower and relatively stable across the older grasslands. Despite the reduction in taxonomic bacterial richness, the functional capacity of the soil microbiome expanded with ecosystem age. Functional richness was lowest in the arable sites but increased significantly within the first 23 years of restoration. Unlike bacterial richness, which dropped and plateaued, functional richness remained elevated in the older grassland soils, suggesting that the communities become functionally more complex over time despite being comprised of fewer bacterial taxa. Concurrently, the composition of the soil community exhibited a monotonic shift toward non-prokaryotic dominance across the chronosequence. The SingleM derived eukaryotic to bacterial gene ratio, utilised as a metric for broad structural change, recorded lowest ratios in the arable soils and significant increases with time. This indicates a continuous proliferation of fungal *and* other non-prokaryotic organisms, with the highest ratios observed in the ancient (143 year) samples.

### Microbial community structure and emergent taxa

Despite the reduction in taxonomic richness, the functional potential of the microbiome expanded over time. Soil genetic richness showed a significant positive response during the 000- to-023-year interval (Figure 1A and 1C). This functional shift was underpinned by the establishment of emergent taxa that were largely absent (<0.01% relative abundance) in arable soils (Figure 2). By year 023, the microbial community was dominated by distinct emergent lineages such as *Aquihabitans sp.* (comprising over 20% of the relative abundance) and members of the Microthrixaceae family. The prominence of Actinobacteria like Candidatus *Microthrix* suggests a functional reorientation toward the decomposition of complex organic matter.

**Figure 2.**
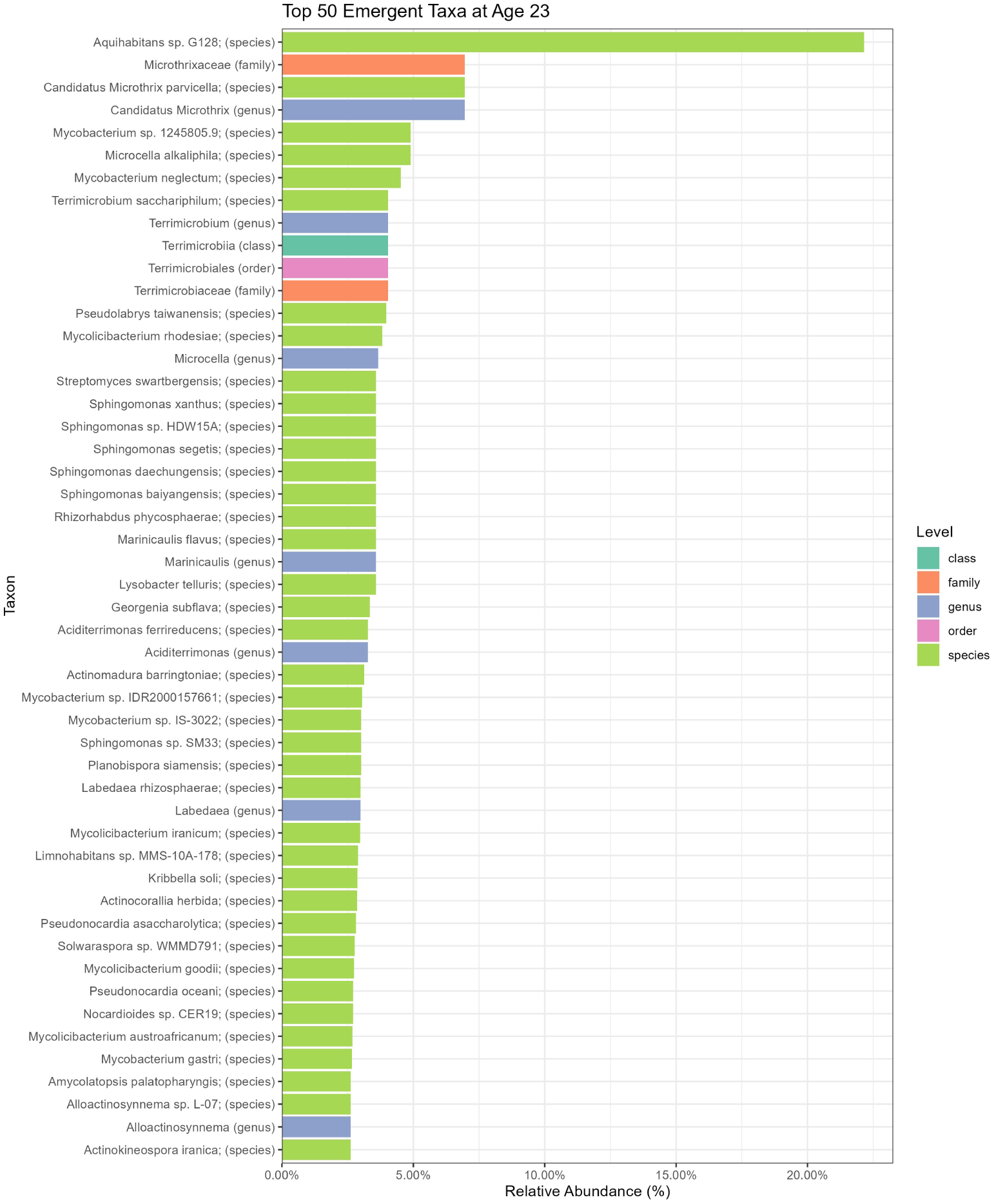
Top 50 emergent microbial taxa during the initial 23-year restoration phase. Taxa were identified via Kaiju protein-level classification across all taxonomic ranks. Emergence criteria were defined as an initial abundance of <0.01% at the 000-year baseline followed by establishment at >0.01% mean relative abundance at the 023-year stage. Taxa are ranked by their proportional representation in the community. Labels indicate the specific taxonomic level for each entry.

### Distinct functional states and metabolic shifts

To understand how the recovery of the soils influenced microbial community functions, we used an ordination method which demonstrated that soil functional composition does not merely recover to a baseline but evolves through distinct, progressively divergent states. This revealed clear clustering based on restoration age, with the first two principal components explaining 18.58% of the total variance (Figure 3). A strong gradient was observed separating the arable and early-restoration (000 and 023 year) sites from the ancient grasslands (143 year). This functional separation was associated with opposing environmental vectors, where arable and early restoration sites were associated with legacies of agricultural improvement (K, P, and Clay content), while ancient sites were defined by vectors associated with ecosystem stability and organic matter sequestration (LOI, Moisture, and C:N ratio). Critically pH, a known master variable did not emerge as a significant driver of functional composition (in-laid table, Figure 3 and Supplementary Figure 2), highlighting the central role of soil organic matter accrual in this system. Notably, the 67-year sites occupied an intermediate ordination space, further confirming that functional equilibrium is not achieved even after six decades of recovery.

**Figure 3.**
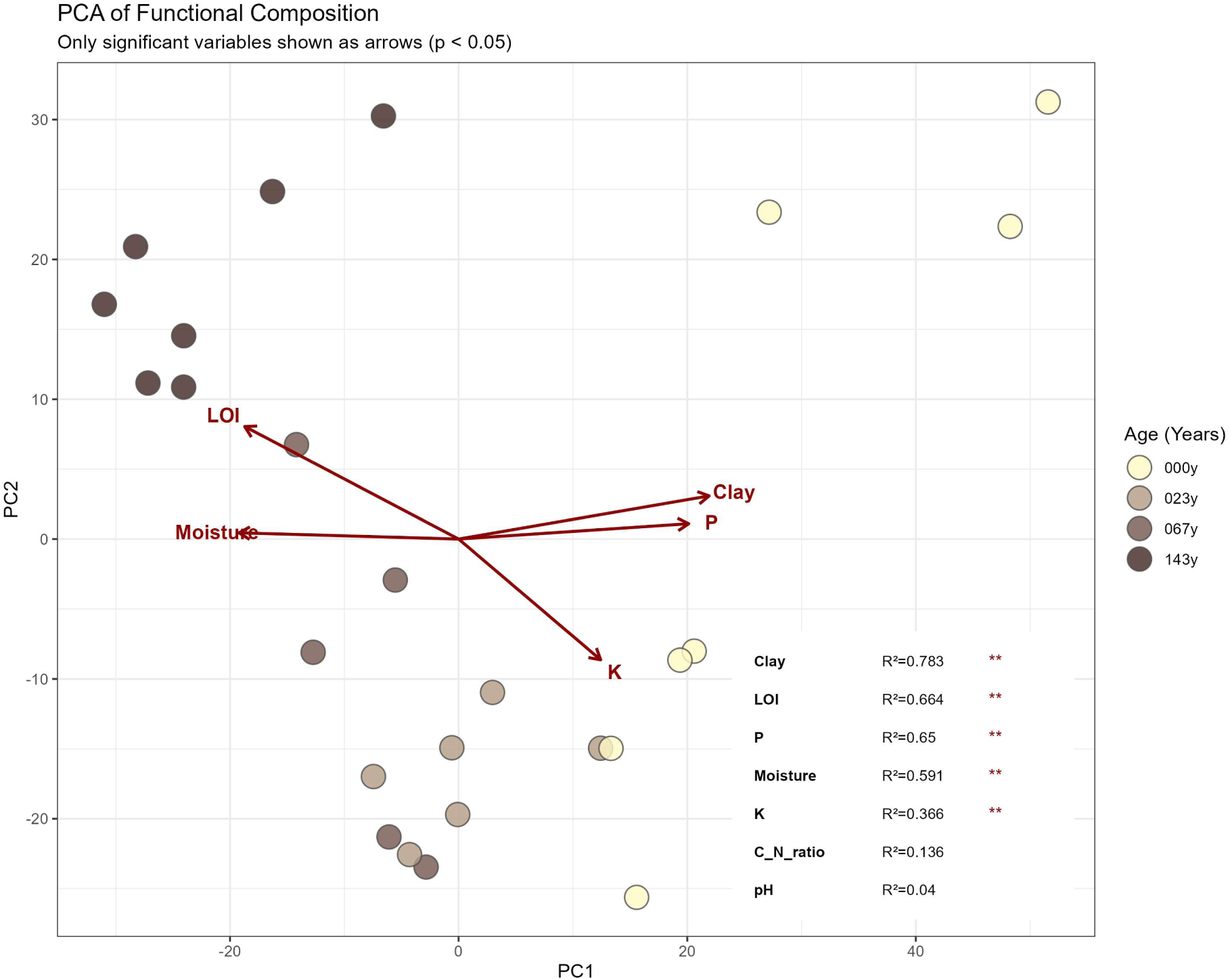
PCA of microbial functional groups with significant environmental drivers. Points represent functional composition, coloured by chronosequence age. Vectors represent significant environmental variables (p < 0.05) based on envfit analysis. Inset table shows goodness-of-fit (R2) and significance codes for all edaphic parameters.

### Emergence and loss of specialised functions and pathways

We identified a suite of functional genes that exhibited monotonic increases or decreases in abundance with restoration age (Figure 4). Restoration of soil led to notable decreases in the abundance of genes associated with membrane transport (ABC-type multidrug transport system), ribosomes (SSU ribosomal proteins), archaeal protein recycling (Archaeal protease subunits), DNA repair (glycosylase), and phage proteins (phage tail proteins). Overall, the recovery of soil condition leads to a reduction in genes associated with growth, maintenance, and disease, along with broader reductions in the subsystems associated with metabolism of proteins, RNA and DNA. Whilst these functions decreased with restoration age, monotonic increases in genes and pathways along the chronosequence were also observed. Subsystems for photosynthesis and motility suggest that soil stability and moisture permit the establishment of mobile, autotrophic or mixotrophic communities. Increases in flagella and aerotaxis sensor receptor proteins enable bacteria to navigate toward optimal oxygen concentrations, maximizing energy production. Fatty acid biosynthesis genes, essential for cell membrane production and membrane fluidity, enable adaptation to environmental stress. Together, these functional gene increases reveal that as soil structural complexity increases with restoration age, soil genetic functions change dramatically. Additionally, the ability to acquire organosulphur compounds becomes more important with restoration age (Alkane sulfonate ABC transporters), suggesting that sulphur availability may be a rate limiting factor. Meanwhile, increases in 4-α-Glucanotransferase suggest greater utilization of complex carbohydrate metabolism, indicating a shift to the utilization of complex substrates as soil organic matter content increases.

**Figure 4.**
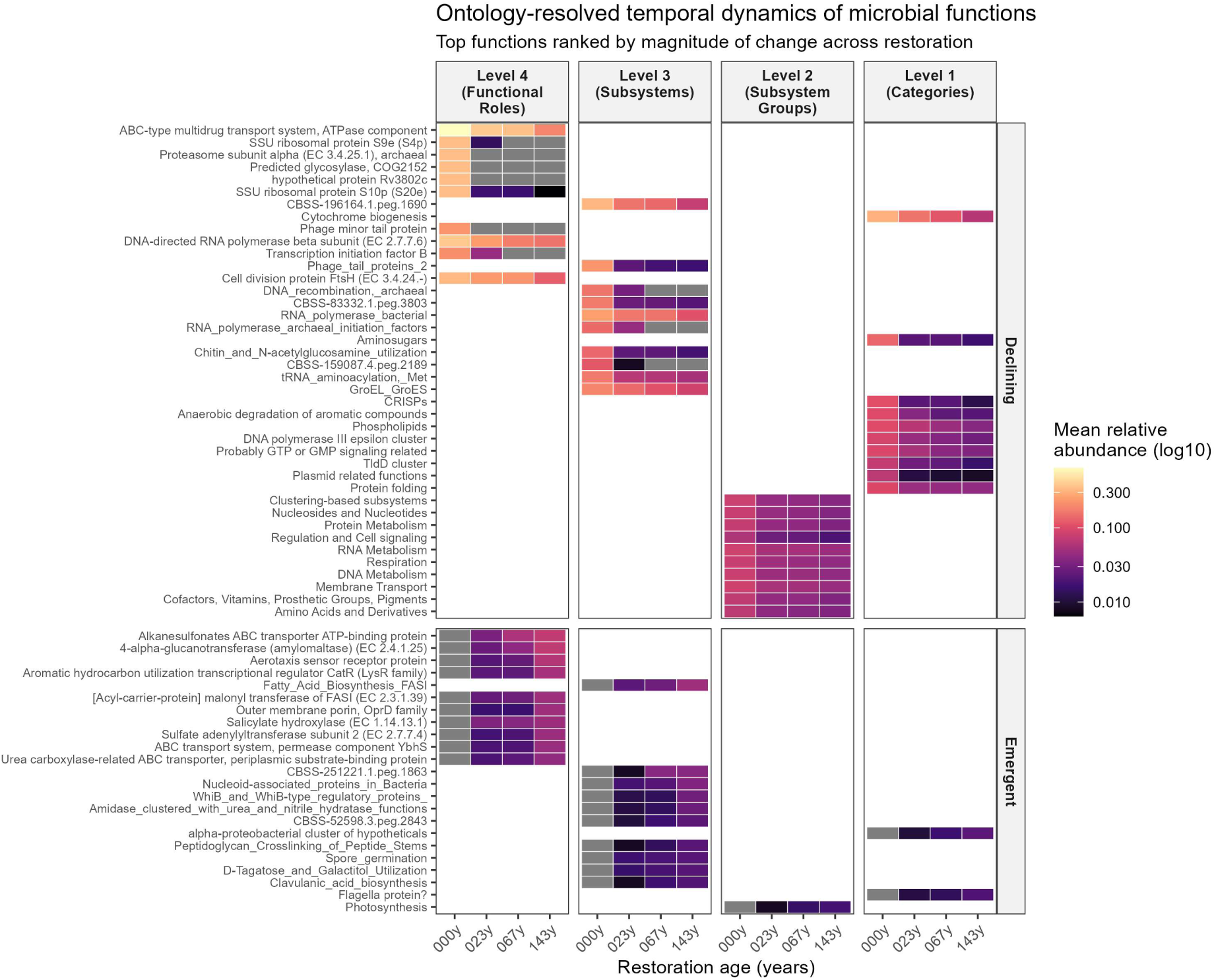
Ontology-resolved temporal dynamics of microbial functional potential across restoration. Heatmap showing mean relative abundances of SEED-annotated microbial functions across restoration age (000, 023, 067, and 143 years). Functions are grouped by SEED hierarchy level (columns) and by direction of change over time (rows), classified as emergent (net increase from 000 to 143 years) or declining (net decrease). Within each hierarchy level and direction, functions are ranked by the magnitude of absolute change in mean relative abundance between 000 and 143 years, and the top functions are displayed. Tile colour indicates mean relative abundance at each time point, shown on a log₁₀-transformed colour scale to enhance contrast among low-abundance functions. This representation highlights both the timing and ontology of functional shifts during ecosystem restoration.

### Genes directly associated with the accumulation of soil organic matter

Soil organic matter (SOM), measured here as LOI, is broadly and consistently identified as a central soil health parameter^33,34^. pH is established as a master variable influencing microbial community composition and function^35^, within this dataset no significant change in pH with restoration age was observed (Figure 3 and Supplementary Figure 2), giving us a unique opportunity to identify and examine genes associated directly with the most important measure of soil health – soil organic matter. Spearman’s rank correlation analysis (Figure 5) identified strong (*rho* > 0.7) inverse relationships between SOM and genes, subsystems, and categories associated with metabolism (proteins, RNA, and DNA), growth (respiration, ribosomes, cell division, and phospholipids), and disease and genetic transfer (CRISPRs, phages, and plasmids). In contrast, strong positive SOM relationships were found with genes for sensing, regulation, and membrane transport (WhiB regulators, transporters, binding proteins, and porins), genes to help exploit greater spatial complexity (flagella, aerotaxis, sporulation, and photosynthesis), genes to help with the utilisation of complex substrates (amylomaltase, hydroxylase, and aromatics), and genes to cope with toxins (clavulanic acid and hydrolase). Notably, increasing SOM led to reduction in many broad categories of Level 1 functions whilst increases in genes with SOM were at the more specific SEED Levels 3 and 4.

**Figure 5.**
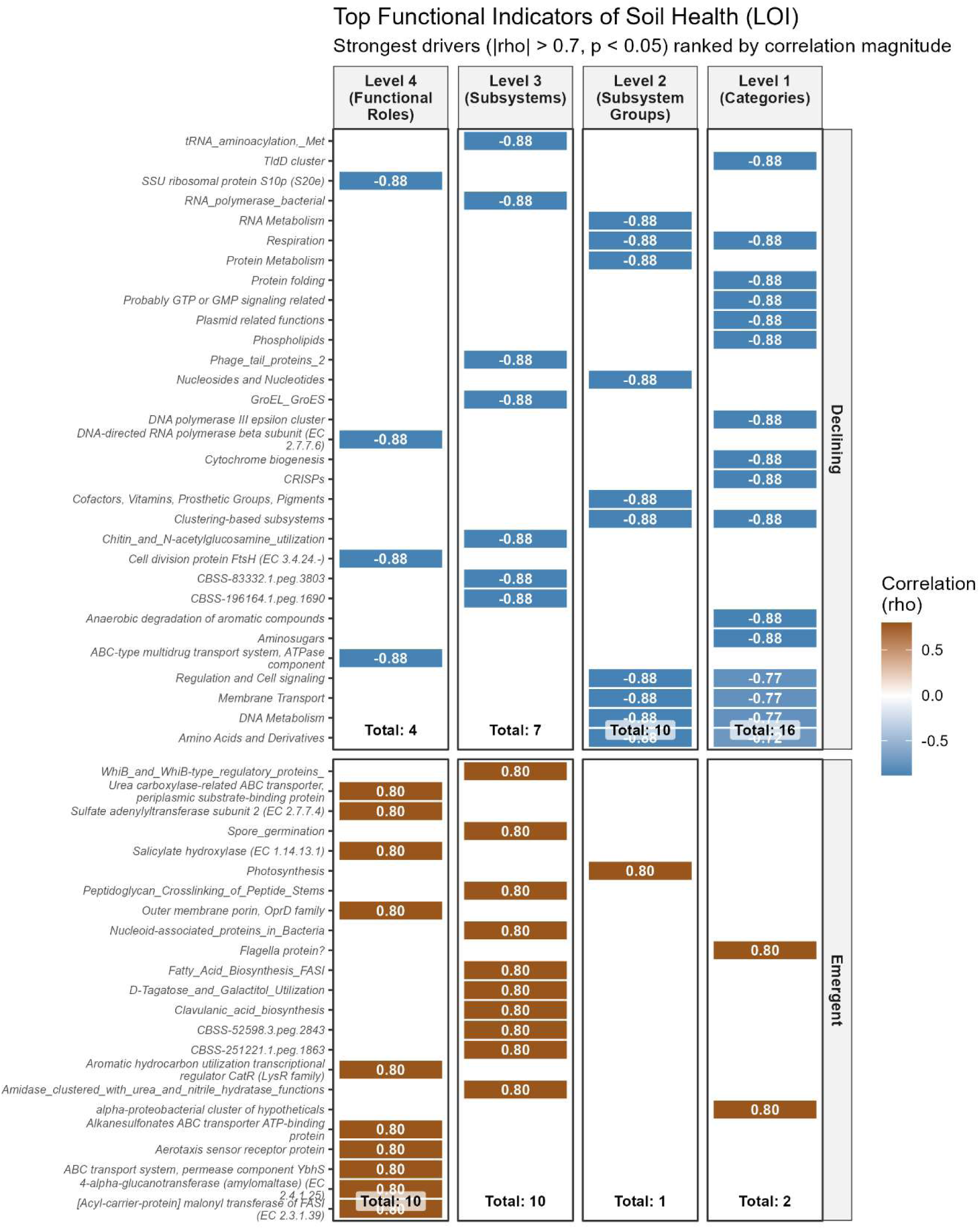
Top functional microbial indicators of soil organic matter (LOI) accumulation across the chronosequence. The heatmap displays microbial functional groups (SEED ontology) that exhibit the strongest statistical coupling with soil organic matter (Loss on Ignition, LOI %). Indicators were selected based on a dual-threshold filter of Spearman’s rank correlation coefficient (rho > 0.7) and significance (p < 0.05) and are strength ordered.

## Discussion

The results of this landscape study challenge the assumption that ecosystem recovery is a synchronous process where below-ground health mirrors above-ground biodiversity. Instead, we observe a complex, multi-phasic recovery where soil properties lag significantly behind vegetation.

### Decoupling of vegetation and soil diversity recovery

The results of our study reveal a fundamental disconnect between the recovery of above-ground floristic diversity and below-ground edaphic function. Congruent with earlier work^10,36–38^, we found that increases in plant diversity and species richness are rapidly achievable. In our study, vegetation richness increased significantly in the first 23 years and continued to rise until before stabilizing, suggesting a saturation of taxonomic diversity within a human lifetime. However, broad diversity metrics can obscure underlying functional deficits. As observed by Redhead et al. (2014)^10^ within this same landscape, even in well-connected landscapes, ancient grasslands show distinct vegetation communities and traits, suggesting that natural regeneration to a full ancient calcareous grassland community takes over a century. While the richness of the vegetation may approach ancient levels relatively quickly, the functional assembly of the plant community remains distinct for much longer. This delay mirrors the persistent legacy effects and slow convergence we observe in the soil ecosystem, which exhibited continued diversification and functional evolution well beyond 67 years. Thus, we observe a two-speed recovery model with a rapid biological reassembly of plant taxonomic richness, followed by a protracted, multi-generational recovery of plant functional traits, soil chemistry, and microbiome architecture.

### The persistence of agricultural legacies and carbon dynamics

The stark contrast in recovery rates is largely associated with the persistence of agricultural legacies^39,40^. Although the visual landscape of the 023-year and 067-year sites resembles ancient grassland, the invisible soil profile remains distinct. The elevated levels of available P and K in the younger restoration sites indicate that the eutrophication caused by historical fertilisation takes decades to attenuate. This nutrient enrichment likely prevents the full establishment of the stress-tolerant, oligotrophic microbial specialists characteristic of ancient soils^41^, and acts as a potential mechanism explaining the observation of that some sites appear to ‘stall’ at a stable, mesotrophic community for long periods^10,39,42^.

Conversely, the accumulation of C and N follow a slow, monotonic trajectory that did not saturate even after 067 years. This suggests that the carbon sequestration potential of calcareous grasslands is not a rapid solution but a long-term service that shows no evidence of saturation within the first 143 years of recovery and may represent a multi-generational process. Consequently, carbon credit schemes or conservation policies based on short-term restoration (<30 years) may severely underestimate the time required to restore the full soil carbon sink in these systems.

### Microbial specialisation reveals efficiency over diversity

A counter-intuitive but critical finding of our study is the inverse relationship between bacterial richness and ecosystem maturity. Arable soils supported the highest bacterial richness^43,44^, likely representing a bloom of fast-growing, disturbance-adapted copiotrophs fuelled by fertiliser inputs and labile carbon^45^. This, in-line with other reports suggesting that arable soils have an increased bacterial diversity. On the other hand, natural regeneration acts as a powerful environmental filter, reducing bacterial taxonomic richness while simultaneously increasing functional richness^46^. This transition likely is consistent with a shift toward more functionally differentiated communities.

This specialisation is further evidenced by the monotonic increase in the non-bacterial-to-bacterial ratio and the emergence of specific taxa such as Microthrixaceae and *Aquihabitans* sp. in the restored sites. The rise of Actinobacteria (like Candidatus *Microthrix*) and fungi suggests a functional reorientation toward the decomposition of more recalcitrant organic matter, consistent with their known ecological roles - a shift that is energetically expensive but essential for the stability of nutrient-poor ancient grasslands.

### Functional maturation and environmental feedback loops

Based on our observations, the restoration of soil function appears to be consistent with plant– soil feedback mechanisms, linking vegetation, soil organic matter, moisture, and the microbiome. This aligns with the concept of plant-soil-feedback (PSF) described by van der Putten et al. (2013)^47^, where the re-establishment of specific plant traits actively conditions the below-ground environment to drive succession. Our analysis of emergent functional genes indicates that as vegetation recovers and contributes root exudates and plant litter, and as soil organic matter builds up to retain moisture, the environment selects for specific metabolic pathways.

In line with this, the enrichment of genes associated with motility and survival in older soils reflects the community’s adaptation to the unbuffered physical stresses and spatial heterogeneity of a natural environment. This contrasts with the homogeneity of managed arable soils, where frequent tillage and inputs buffer microorganisms against physico-chemical variation. This trajectory mirrors the trade-off described by Fierer et al. (2007)^48^ and Ho et al. (2017)^49^, who noted a shift from r-strategist (copiotrophic) dominance in disturbed, nutrient-rich soils to K-strategist (oligotrophic/stress-tolerant) dominance in stable, late-successional ecosystems. Notably, soil pH, widely regarded as the ‘master variable’ determining global soil microbial community structure ^43,44^, did not change in this system, usefully decoupling it as a driver of observed functional change.

### Lack of equilibrium and conservation implications

Importantly, the most significant implication of this study is the apparent lack of a terminal equilibrium within the human management timeframe. The ordinations and rate-of-change analyses demonstrated that 067-year-old grasslands are functionally distinct from 143-year-old ancient sites. The trajectory of change, particularly in C, moisture, and Mg, suggest that restoration is a continuous process spanning multiple human generations^50^. This challenges binary definitions of restored versus degraded land^51^. Therefore, conservation policy must recognise that 60-year-old regeneration sites are valuable but transitional ecosystems, requiring continued protection to allow the slow accumulation of the functional complexity, specifically the fungal networks and specialised metabolic depth, which characterises these ancient soils.

### Conclusion and future research

Looking forward, this 160-year chronosequence necessitates a paradigm shift in how we value and manage restoring landscapes, moving beyond the assumption that ecosystem recovery is a rapid or synchronous process. The persistence of agricultural nutrient legacies and the lack of functional convergence even after seven decades demonstrate that current conservation frameworks, often bound by short-term funding cycles, are ill-equipped to capture the full trajectory of edaphic recovery. To address the temporal gap between rapid vegetation recovery and slower soil functional maturation, restoration strategies may benefit from considering approaches that extend beyond passive regeneration. Our results suggest that dispersal limitation and persistent nutrient legacies could constrain the re-assembly of specialised microbial communities. One potential accelerant is targeted soil inoculation; whereby early restoration sites receive small amounts of donor soil from long-established grasslands to reintroduce oligotrophic taxa such as *Aquihabitans* and members of the *Microthrixaceae*. Experimental studies have demonstrated that whole-soil inoculation can help overcome dispersal barriers and steer plant and soil community assembly toward reference states, accelerating the development of late-successional functions^52^. While not directly tested here, such approaches may enhance the recovery of tightly coupled nutrient cycling processes that often remain incomplete after decades of spontaneous regeneration^53^.

The persistence of elevated phosphorus in younger sites suggests that nutrient legacies may constrain microbial reassembly, indicating that restoration management could benefit from exploring targeted nutrient reduction strategies aimed at gradually re-establishing higher C:N ratios. Such stoichiometric shifts may favour the development of more stable and functionally complex microbial communities. In addition, our findings highlight a potential mismatch between rapid vegetation recovery and slower soil functional maturation, suggesting that monitoring frameworks may benefit from incorporating below-ground indicators alongside above-ground metrics. For example, metagenomic assessment of genes associated with stress tolerance and carbon cycling could provide complementary indicators of functional development. More broadly, the continued increase in soil organic matter across the chronosequence suggests that soil carbon accrual operates over multi-decadal to centennial timescales, reinforcing the importance of aligning restoration expectations with the long temporal dynamics of soil ecosystem recovery.

---

**Supplementary Figure 1.**
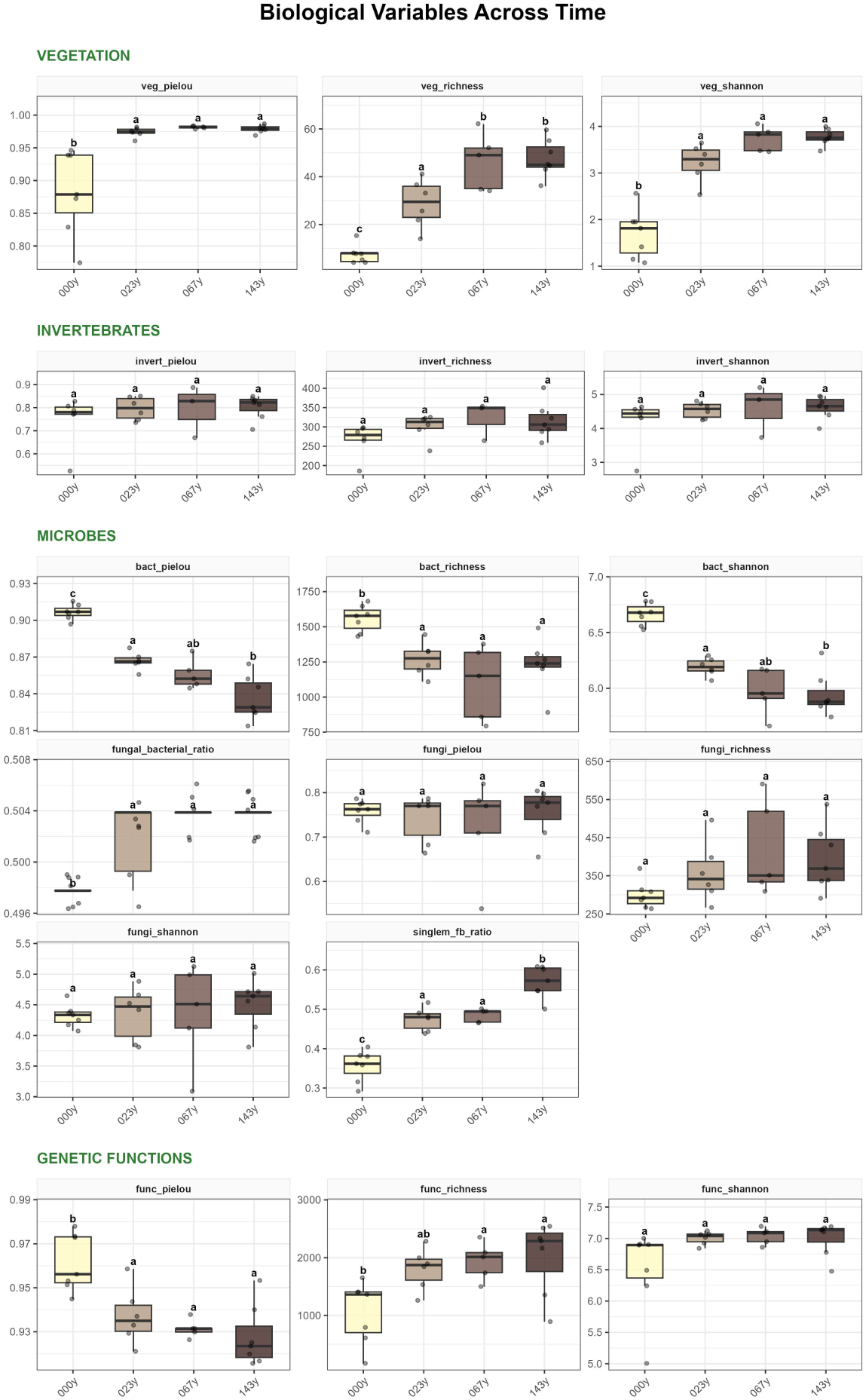
Successional dynamics of biological variables across a 143-year soil chronosequence. Box-and-whisker plots represent the distribution of vegetation (richness, Shannon diversity, and Pielou’s evenness), invertebrate communities, microbial indices (including bacterial and fungal metrics, PLFA derived fungal-bacterial ratio and SingleM derived eukaryote-bacterial gene ratio), and genetic functions. Data points are jittered to show individual sample distribution. The colour gradient represents soil age, transitioning from light yellow (000y) to dark brown (143y). Different lowercase letters above boxes indicate significant differences between age groups based on Tukey’s Honestly Significant Difference (HSD) test (p < 0.05).

**Supplementary Figure 2.**
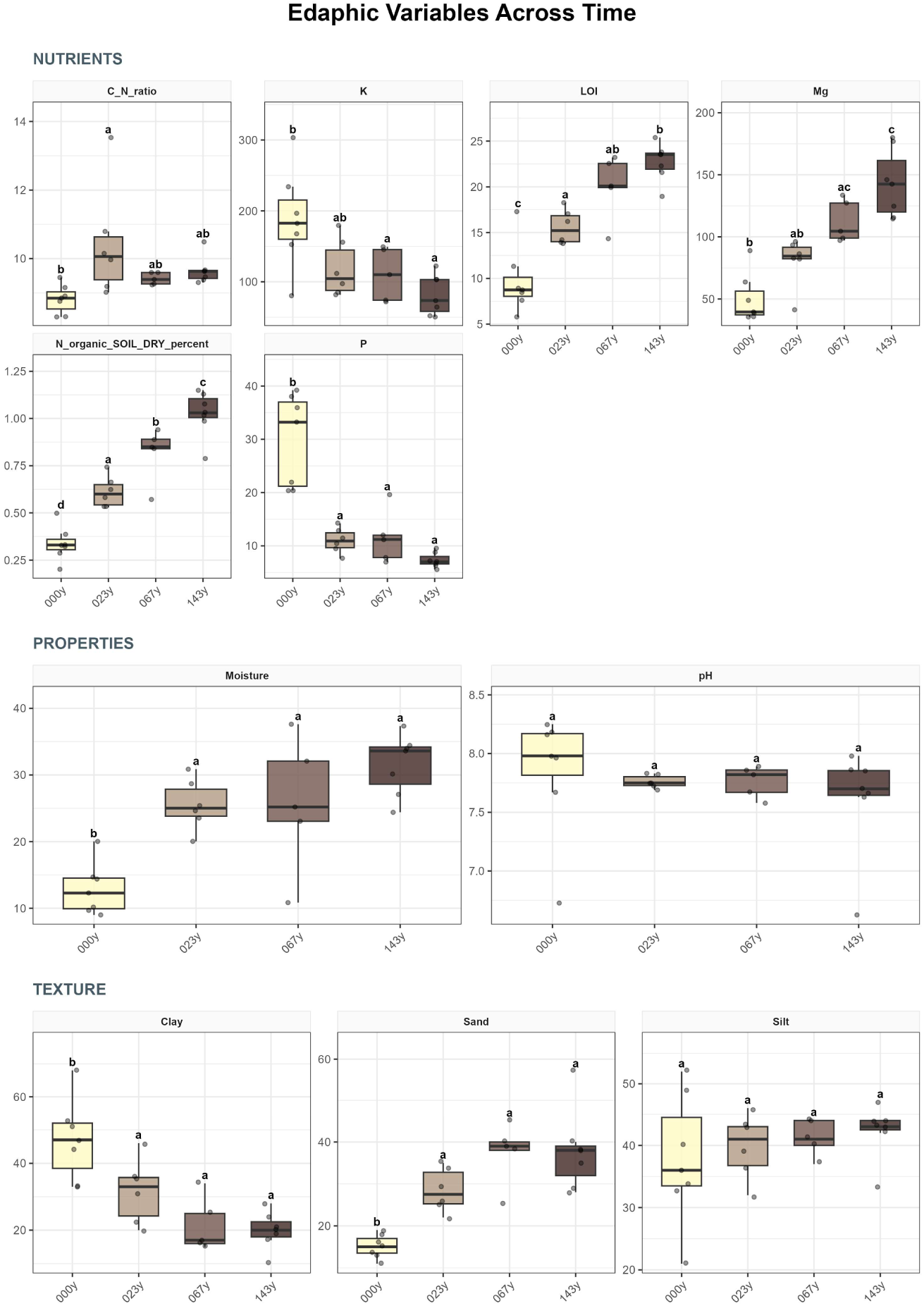
Evolution of edaphic properties and soil texture over 143 years of soil development. Panels illustrate changes in soil nutrients (LOI, organic Nitrogen, C:N ratio, Phosphorus, Potassium, and Magnesium), physical properties (Moisture and pH), and soil texture (Sand, Silt, and Clay content). The colour ramp follows the chronosequence from light yellow (000y) to dark brown (143y). Boxplots display the median and interquartile range; whiskers extend to the furthest non-outlier data point. Shared lowercase letters indicate no significant difference between time points according to Tukey’s HSD post-hoc analysis (p > 0.05).

